# ModEst - Precise estimation of genome size from NGS data

**DOI:** 10.1101/2021.05.18.444645

**Authors:** Markus Pfenninger, Philipp Schönnenbeck, Tilman Schell

## Abstract

Precise estimates of genome sizes are important parameters for both theoretical and practical biodiversity genomics. We present here a fast, easy-to-implement and precise method to estimate genome size from the number of bases sequenced and the mean sequence coverage. To estimate the latter, we take advantage of the fact that a precise estimation of the Poisson distribution parameter lambda is possible from truncated data, restricted to the part of the coverage distribution representing the true underlying distribution. With simulations we could show that reasonable genome size estimates can be gained even from low-coverage (10X), highly discontinuous genome drafts. Comparison of estimates from a wide range of taxa and sequencing strategies with flow-cytometry estimates of the same individuals showed a very good fit and suggested that both methods yield comparable, interchangeable results.

## Introduction

Eukaryotic genomes vary tremendously in size (Oliver *et al*. 2007; Bennett & Leitch 2005; Petrov 2001; Kapusta *et al*. 2017; Carta *et al*. 2020), yet the underlying processes for this variability are not yet fully understood (Elliott & Gregory 2015). To understand and study mechanisms of genome size variation, such as proliferation of repetitive elements (Blommaert *et al*. 2019), effective population size (Lefébure *et al*. 2017; Lynch & Conery 2003) or correlation to other traits (Gardner *et al*. 2020; Prokopowich *et al*. 2003), reliable estimates for the taxon under scrutiny are therefore mandatory. This is all the more important as substantial changes in genome size may even occur among closely related sister species, i.e. over relatively short evolutionary time scales (Keyl 1965; Agudo *et al*. 2019, Vitales et al. 2020). A precise estimation of genome size is also important for genomic projects. For example, in the assembly of genomes, the proportion of the true genome size covered by a given assembly draft is a quality criterion and limits the maximum size of the draft. Also resequencing projects requiring a certain coverage e.g. for genotyping profit from a reliable genome size estimate (Fountain *et al*. 2016).

Flow cytometry is generally deemed to yield reliable estimates of genome size (Johnston *et al*. 2019; Doležel & Greilhuber 2010). Yet, this method is not without caveats (Wang *et al*. 2015) and requires specialised laboratory skills and availability of the relatively expensive equipment. Moreover, the method depends on availability of fresh or frozen tissue with largely intact cells, which narrows the range of taxa for which such analyses are practically feasible (Johnston *et al*. 2019).

Bioinformatical analysis of next generation sequencing data provides an alternative for estimating genome size (Vurture *et al*. 2017). Besides the widely used k-mer based methods (Lipovský *et al*. 2017; Li & Waterman 2003), Schell et al. 2017 introduced a very simple method for genome size estimation, relying on mapping statistics of NGS reads mapped back to a draft assembly. The approach assumes that the probability to sequence a genome position is identical over the entire genome, i.e. that their true coverage is Poisson distributed. Even though there is a slight bias regarding the double strand breaking positions during DNA preparation for NGS sequencing, the impact on the resulting sequencing coverage distribution is negligible (Poptsova et al. 2014). In a perfect assembly covering the entire genome, lambda as the parameter of the underlying Poisson distribution (as well as the mean and median) of the coverage distribution should therefore be identical to the true coverage. Dividing the number of sequenced, successfully back-mapped bases by the lambda of the observed coverage should yield a precise estimate of the true genome size. In most real draft genomes, however, repetitive regions are not resolved which results in collapsed repeat regions, and in an assembly that is shorter than the true length (Treangen & Salzberg 2012). These collapsed repeat regions are over-proportionally covered, skewing the coverage distribution, and hence, estimates of lambda upwards. A second source of systematic error in assemblies are relatively diverged heterozygous regions, e.g. from inversions that are not identified as homologous. These will result in a double representation of the respective region in the genome, making it longer (Asalone *et al*. 2020). Consequently, the expected coverage of these regions in the assembly will be half of the true coverage and skew the coverage distribution and parameters estimated from it downwards. In real genome assemblies, both errors likely occur to various extents (Sohn & Nam 2018), rendering a naïve use of parameters estimated from the observed coverage distribution misleading.

We show here how the observed coverage distribution and an estimate of the number of bases sequenced from genome assembly drafts can be used to infer precise estimates of genome size. We name the approach ModEst from **Mod**al **Est**imation of genome size. We tested the methods with simulations, including various degrees of divergent heterozygous sites and a tetrapoid genome, and compare genome size estimates from real data over a wide range of genome sizes with those derived from flow cytometry and k-mer based methods.

## Material and Methods

### Theoretical background

Under the assumption that NGS sequencing methods sequence all bases in a genome with equal probability, dividing the number of bases sequenced (*N*) by the true length of the genome (*L*) yields the mean or expected coverage (*c*) (Sims *et al*. 2014).

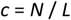

Since the coverage distribution is discrete, it can be modelled by a Poisson distribution with parameter λ as *c*. As we are interested in *L*, we need to find reliable estimates for *N* and *c* from empirical data.

The number of bases used for the assembly of a particular genome is usually known. This number is, however, not necessarily identical to the number of bases sequenced from the target genome. Depending on the origin of the DNA, the data set may contain more or less reads originating from contaminations, the microbiome, and certainly reads from the mitochondrial or plastid genomes (Kumar *et al*. 2013). Even though several tools and pipelines exist to remove the bulk of such reads (Chaliis et al. 2020), this rarely succeeds completely. The number of bases after thorough cleaning, *N*_*clean*_, estimates therefore rather the upper limit of *N*.

An alternative is the number of bases mapped back to the genome assembly draft *N*_*bm*_. For this number to represent a good approximation of the number of bases sequenced from the corresponding genome, all genomic elements (telomers, centromers, repeats) must be represented in the assembly at least once without presence of contamination etc. and all reads must map back. This number is therefore a lower limit estimator of *N*.

As detailed in the introduction, the empirical coverage distribution of back-mapped reads is usually biased by errors in the genome draft due to collapsed repeats and/or other assembly errors. However, commonly at least a substantial part of the back-mapped reads map to unique sequences in the genome draft and should consequently show a coverage distribution following the true underlying Poisson distribution. Estimating *λ* from the part of the distribution we know is not biased by assembly errors should therefore yield a reliable estimator of *c*. In Schell et al. 2017, the modal value of the empirical coverage distribution (*m*), i.e. the most often observed coverage was used as an estimator of *c*. The modal value is a fairly good approximation of *λ* because the difference is in all cases smaller than or equal to 1 and therefore becomes relatively less biased when *λ* is high (i.e. high mean coverage). Nevertheless, better methods for estimating *λ* from truncated Poisson distributions exist (Delignette-Muller & Dutang 2015; Nadarajah & Kotz 2006; Böhning & Schön 2005; David & Johnson 1952).

As mentioned above, the coverage distribution may show more than a single peak. One possibility to obtain a bimodal distribution arises from highly divergent heterozygous tracts in the respective genome. In the assembly process, such divergent tracts may not be identified as homologous by the algorithm and thus occur as separate regions. Consequently, the coverage in such areas is only half the true coverage. If a considerable proportion of the genome consists of such divergent heterozygous regions, a second peak may appear in the coverage histogram. It has its maximum usually at half the coverage of the larger peak. In this case, the peak with the larger coverage represents the true coverage. Except for recent hybrid individuals, the latter peak should nevertheless always be the higher one.

Another possibility to obtain a multimodal coverage distribution arises from polypoid species. If the multiplied genomes diverged to an extent that both are completely represented in the assembly, the genome size estimation process is not any different from a diploid species. The other extreme would be a multiplied genome that is so little diverged that only a single copy appears in the assembly. In an intermediate stage, some more diverged parts of the multiplied genomes may be resolved, while others are collapsed in the assembly. The collapsed parts are expected to be over-covered and therefore the lowest peak represents the true coverage.

In general, the observation of a multimodal coverage distribution of the backmapped reads is indicative of issues with the assembly. Genome size estimation with the proposed ModEst method should be nevertheless possible, given appropriate caution.

### Practical approach

All the figures needed to estimate the genome size according to the method described here are usually collected in the process of genome assembly or can be easily calculated with standard tools. In particular, samtools stats and bedtools genomecov can be used for this purpose. The output of samtools stats provides information on bases sequenced and mapped, while the output of bedtools genomecov provides the empirical coverage distribution. The latter can be used as input for R. After preparing the data, we first estimated the modal value of the empirical distribution. This modal value is used as starting point for a Maximum Likelihood method to estimate *λ* from a truncated Poisson distribution as implemented in the R-libraries *truncdist* and *fitdistrplus* (Delignette-Muller & Dutang 2015; Nadarajah & Kotz 2006). We empirically determined suitable upper and lower truncation limits and give recommendations below. The respective R-code can be found in the Supplement and a Perl wrapper-script, including all necessary dependencies can be found at https://github.com/schellt/backmap.

### Simulations

To illustrate the influence of factors like sequencing depth, genome size, repeat content and - distribution on the different genome size estimation methods, we simulated five different genomes according to real examples. Publicly available genome assemblies and annotations of *Saccharomyces cerevisae, Caenorhabditis elegans, Arabidopsis thaliana, Drosophila melanogaster* and *Scophthalmus maximus* were used to obtain distributions of size and distance between annotated repeat regions. Simulated genomes of the size of the five genome assemblies mentioned above were then created using a custom Python-tool, available at https://github.com/Croxa/Simulate-Genome. Regions annotated as repeat regions (rr) were filled with random repeat units up to 10 bp length, high complexity regions with random nucleotides. For sake of ease, we simulated the genomes on a single chromosome. A mean GC content of 0.5 was applied to both categories. Characteristics of the simulated genomes can be found in Table 1.

**Table 1:** Simulated genomes and their characteristics, rr = repeat regions.

| Simulated genome | Size<br>(Mbp) | average count<br>of bases<br>between rr | average count<br>of bases of rr | % of rr |
| --- | --- | --- | --- | --- |
| 1 *Saccharomyces cerevisiae*-like | 12 | 1246.68 | 156.67 | 5.26 |
| 2 *Caenorhabditis elegans*-like | 100 | 508.66 | 166.42 | 13.23 |
| 3 *Arabidopsis thaliana*-like | 120 | 622.32 | 311.55 | 18.06 |
| 4 *Drosophila melanogaster*-like | 144 | 372.42 | 242.48 | 23.39 |
| 5 *Scophthalmus maximus*-like | 524 | 521.84 | 45.64 | 3.74 |

From these simulated genomes, we generated synthetic next-generation sequencing short read sets of 10X, 30X and 60X coverage using ART Illumina 2.5.8 (Huang *et al*. 2012). This tool emulates the sequencing process with built-in, technology-specific read error models, base quality value profiles parameterized empirically for large sequencing datasets and even adds the sequencing adapters. The reads were simulated paired-end, length of 150 bp with a standard deviation of 10 and an insert size of 300 bp. The Illumina sequencing system profile was HiSeq 2500 (HS25).

The read sets were trimmed with Trimmomatic 0.39 (Bolger *et al*. 2014). Trimmed were usual Illumina adapters (ILLUMINACLIP:adapter.fa:2:30:10), leading and trailing bases with a quality score lower than 5, sliding windows with the size of 20 and an average quality score below 5 and reads with a length of 50 or lower.

In a first set of experiments, the trimmed read sets of different coverage were back-mapped to the simulated genomes they were derived from. Mapping was executed within the wrapper script backmap.pl using bwa mem 0.7.17 without changing default options from backmap.pl. BWA (Burrows-Wheeler Aligner) is a widely used algorithm for mapping low-divergent sequences against a large reference genome (Li 2013).

To estimate the influence of genome assemblies of varying quality on the accuracy of the genome size estimate, we assembled each read set with SPAdes, the St. Petersburg genome assembler. This algorithm is implemented in a toolkit containing various assembly pipelines (Bankevich *et al*. 2012). SPAdes 3.13.0 was used to assemble both trimmed paired and unpaired reads in a one-pass assembly using default options. The respective read sets were back-mapped and analysed as described above. For one simulation (*A. thaliana*-like, 10X coverage), we evaluated the effect of different truncation limits on the precision of the *λ* estimation. For coverage class windows ranging from 11 to 5, centred on the modal value, the deviation of the ML estimate decreased from 0.4% to 4%. We performed the *λ* calculations therefore with a window size of eleven around the estimated modal value.

The influence of different amounts of diverged heterozygous genome stretches on size estimation was evaluated using the Saccharomyces-like genome. We simulated the genome with X,Y and Z% heterozygous stretches. To make sure that these stretches were not collapsed in the assembly process, we chose a sequence divergence of 10%. Likewise, we inferred the effect of polyploidy on genome size estimation with our method. We doubled the Saccharomyces-like genome and randomly changed bases in the complex part of one of the genomes. We simulated divergences of 0.5%, 1% and 5% among the two genomes. Both sets of simulations were performed as described above with 30X coverage.

For all simulations, we calculated four different genome size estimates:

i. N_clean_/*λ*, the number of “sequenced” bases after cleaning and trimming divided by the truncated Poisson ML *λ* estimate derived from the empirical coverage distribution.
ii. N_clean_/*m*, the number of “sequenced” bases after cleaning and trimming divided by the modal value of the empirical coverage distribution.
iii. N_bm_/*λ*, the number of back-mapped bases divided by the ML *λ* estimate derived from the empirical coverage distribution.
iv. N_bm_/*m*, the number of back-mapped bases divided by the modal value of the empirical coverage distribution.

For each estimate, we calculated the relative deviation from the true known genome size.

### Empirical data

We used data from de novo genome assemblies that were sequenced in the last few years at the LOEWE Translational Biodiversity Genomics Centre and for which flow cytometry estimates from the same individual/clone/population were available. The taxonomic range of genomes comprised plants and several animal taxa with a focus on insects (Table 2).

**Table 2.** Genomes used for empirical evaluation.

| Species | Taxon | Flow-cytometry estimate [Mb] | Backmapping estimate [Mb] | k-mer based estimate [Mb] | Sequencing technique | Citation |
| --- | --- | --- | --- | --- | --- | --- |
| *Hydropsyche tenuis* | Insecta | 260.6 | 228.6 | 222.8 | Short read | Heckenhauer *et al.* 2019 |
| *Plectrocnemia conspersa* | Insecta | 455.2 | 364.9 | 316.3 | Short read | Heckenhauer *et al.* 2019 |
| *Agapetus fuscipens* | Insecta | 721.8 | 583.5 | 463.2 | Short read | Heckenhauer *et al.* 2021 |
| *Odontocerum albicorne* | Insecta | 1616.0 | 1270.0 | 1103.4 | Short read | Heckenhauer *et al.* 2021 |
| *Drusus annulatus* | Insecta | 840.2 | 684.3 | 592.3 | Short read | Heckenhauer *et al.* 2021 |
| *Halesus radiatus* | Insecta | 1212.4 | 972.3 | 918.7 | Short read | Heckenhauer *et al.* 2021 |
| *Micropterna sequax* | Insecta | 1434.7 | 1100.0 | 981.7 | Short read | Heckenhauer *et al.* 2021 |
| *Micrasema longulum ML1* | Insecta | 663.6 | 707.7 | 650.7 | Short read | Heckenhauer *et al.* 2021 |
| *Micrasema longulum ML3* | Insecta | 663.6 | 637.8 | 635.2 | Short read | Heckenhauer *et al.* 2021 |
| *Micrasema minimum* | Insecta | 588.8 | 329.3 | 333.8 | Short read | Heckenhauer *et al.* 2021 |
| *Rhyacophila evoluta Rss1* | Insecta | 651.3 | 581.8 | 518.8 | Short read | Heckenhauer *et al.* 2021 |
| *Rhyacophila evoluta HR1* | Insecta | 651.3 | 565.5 | 514.4 | Short read | Heckenhauer *et al.* 2021 |
| *Glaux maritima* (also known as *Lysimachia maritima*) | Angiosperm plant | 1270.0 | 1541.4 | 1221.3 | Short read | Segers *et al.* unpublished data |
| *Radix auricularia* | Mollusca | 1575.0 | 1603.0 | 947.1 | Short read | Schell *et al.* 2017 |
| *Crematogaster levior* | Insecta | 455.0 | 356.0 | 255.9 | Short read | Hartke *et al.* 2019 |
| *Daphnia galeata* | Crustacea | 155.0 | 157.0 | 150.5 | Short read | Nickel *et al.* 2021 |
| *Candidula unifasciata* | Mollusca | 1540.0 | 1420.0 | 977.6 | Short read | Chueca *et al.* 2021a |
| *Styela plicata* | Tunicata | 430.9 | 468.6 | 338.8 | Short read | Galià-Camps *et al.*<br>unpublished data |
| *Callionymus lyra* | Teleostei | 645.0 | 653.2 | 562.0 | Short read | Winter *et al.* 2020 |
| *Pimpla turbinella* | Insecta | 300.0 | 298.0 | 206.0 | Short read | Reumont *et al.*<br>unpublished data |
| *Fagus sylvatica* | Angiosperm<br>plant | 582.4 | 542.0 | 541.0 | Short read | Mishra *et al.* 2021 |
| *Aedes japonicus* | Insecta | 857.0 | 836.3 | 699.0 | Short read | Reuss *et al.*<br>unpublished data |
| *Nyctereutes procynoides* | Mammalia | 3100.0 | 3230.0 | - | Long read | Chueca *et al.* 2021b |
| *Microthlaspi erraticum* | Angiosperm<br>plant | 194.5 | 211.0 | 211.00 | Short read | Mishra *et al.* 2020 |
| *Crematogaster levior*,<br>*species B* | Insecta | 390.0 | 406.7 | - | Long read | Feldmeyer *et al.*<br>unpublished data |
| *Camponotus femoratus* | Insecta | 330.0 | 340.0 | - | Long read | Feldmeyer *et al.*<br>unpublished data |
| *Astacus astacus* | Crustacea | 16891.0 | 16750.0 | - | Short read | Theissingner *et al.*<br>unpublished data |
| *Lamprophis fuliginosis* | Squamata | 1480.0 | 1617.0 | - | Long read | Hiller *et al.*<br>unpublished data |
| *Desmodus* | Mammalia | 2337 | 2089 | - | Long read | Hiller *et al.*<br>unpublished data |

If not stated otherwise in the citations, genome size estimates from flow cytometry were estimated following a protocol with propidium iodide-stained nuclei described in (Hare & Johnston 2012). Tissue of the organism was chopped with a razor blade in a petri dish containing 2 ml of ice-cold Galbraith buffer. The suspension was filtered through a 42-μm nylon mesh and stained with the intercalating fluorochrome propidium iodide (PI, Thermo Fisher Scientific) and treated with RNase II A (Sigma-Aldrich), each with a final concentration of 25 μg/ml. The mean red PI fluorescence signal of stained nuclei was quantified using a Beckman-Coulter CytoFLEX flow cytometer with a solid-state laser emitting at 488 nm. Fluorescence intensities of 5000 nuclei per sample were recorded. We used the software CytExpert 2.3 for histogram analyses The total quantity of DNA in the sample was calculated as the ratio of the mean red fluorescence signal of the 2C peak of the stained nuclei of the target organism divided by the mean fluorescence signal of the 2C peak of the reference standard times the 1C amount of DNA in the standard reference. Six replicates were measured on six different days to minimize possible random instrumental errors. We report the mean value of these measurements.

For each of the genomes, we calculated N_bm_/*m* since we could not reconstruct the exact state of taxonomic read cleaning i.e. removal of contamination reads from other taxa for all genomes. The modal value was chosen, because the coverage exceeded 50X in most cases. For comparison, we performed or used published k-mer based estimates as far as available. First a k-mer profile was generated from Illumina reads using jellyfish 2.3.0 tools (Marçais & Kingsford 2011) count with a length of *k=*21 and counting k-mers on both strands and histo. Subsequently, the generation histogram was used as input for the GenomeScope webserver (Vurture *et al*. 2017) together with the above mentioned length of k and read length. For some organisms, the approach could find no appropriate model. In addition, it is not suitable for long read technologies.

### Statistical analysis

The performance of the two bioinformatic genome size estimation methods was evaluated by their linear regression fit with the respective flow-cytometry estimates. We compared the two slopes of the regression for statistical difference (Cohen *et al*. 2013).

## Results

### Simulations

The single-pass assemblies derived from the simulated short reads were highly fragmented with thousands of short scaffolds, almost independent of simulated coverage (Table 3). For the *S. saccharomyces*-like, the *C. elegans*-like and the *S. maximus*-like genomes, the total lengths of the assemblies were above 90% of the true size, for the remaining two below 80%. This was reflected in the back-mapping rates that were highly correlated to the relative assembly length (r = 0.995, p < 0.001, Table 3).

**Table 3.** Characteristics of simulated genomes, their assemblies, back-mapping and estimation of the parameter of the underlying Poisson-distribution.

| Simulated genome | simulated coverage | true size [Mbp] | assembly size [Mbp] | proportion of true length | number of contigs | mean contig length [bp] | Mbp "sequenced" =N<sub>clean</sub> | bp mapped = N<sub>m</sub> | proportion of bases mapped |
| --- | --- | --- | --- | --- | --- | --- | --- | --- | --- |
| *Saccharomyces_cerevisae*_like | 10 | 12.08 | 11.24 | 0.930 | 2,021 | 5,561 | 120.8 | 113.0 | 0.931 |
|  | 30 | 12.08 | 11.24 | 0.931 | 1,823 | 6,170 | 362.4 | 339.1 | 0.932 |
|  | 60 | 12.08 | 11.24 | 0.932 | 1,906 | 5,905 | 724.8E+08 | 677.7 | 0.931 |
| *Caenorhabditis_elegans*_like | 10 | 100.0 | 92.77 | 0.927 | 105,104 | 883 | 100.0E+09 | 933.7 | 0.904 |
|  | 30 | 100.0 | 9280 | 0.928 | 98,341 | 944 | 300.1E+09 | 2807 | 0.910 |
|  | 60 | 100.0 | 92.25 | 0.929 | 99,957 | 930 | 600.2E+09 | 5598 | 0.904 |
| *Arabidopsis_thaliana*_like | 10 | 120.1 | 92.83 | 0.773 | 66,881 | 1,388 | 1201 | 956.6 | 0.780 |
|  | 30 | 120.1 | 93.03 | 0.775 | 63,695 | 1,461 | 3602 | 2881 | 0.785 |
|  | 60 | 120.1 | 92.75 | 0.772 | 61,615 | 1,505 | 7205 | 5759 | 0.784 |
| *Drosophila_melanogaster*_like | 10 | 144.1 | 107.8 | 0.748 | 104,002 | 1,037 | 1441 | 1118 | 0.755 |
|  | 30 | 144.1 | 107.7 | 0.747 | 95,701 | 1,125 | 4322 | 3382 | 0.763 |
|  | 60 | 144.1 | 107.6 | 0.747 | 95,523 | 1,127 | 8643 | 6725 | 0.756 |
| *Scophthalmus_maximus*_like | 10 | 524.1 | 523.4 | 0.999 | 76,507 | 6,842 | 5241 | 5220 | 0.994 |
|  | 30 | 524.1 | 524.1 | 1.000 | 63,360 | 8,261 | 15720 | 15670 | 0.995 |
|  | 60 | 524.1 | 425.1 | 1.000 | 63,260 | 8,274 | 31440 | 31340 | 0.995 |
| *Saccharomyces_cerevisae*_like 1% divergent heterozygous regions | 30 | 12.08 | 11.78 | 0.975 | 1,152 | 10,226 | 356.1 | 352.2 | 0,989 |
| 5% divergent heterozygous regions | 30 | 12.08 | 12.18 | 1.008 | 3,020 | 4,032 | 354.0 | 350.5 | 0,990 |
| 10% divergent heterozygous regions | 30 | 12.08 | 12.68 | 1.049 | 5,435 | 2,333 | 351.4 | 347.1 | 0,988 |
| 10% divergent heterozygous regions | 30 | 12.08 | 13.68 | 1.133 | 10,163 | 1,346 | 346.1 | 341.9 | 0,988 |
| Tetraploid *Saccharomyces_cerevisae*_like 0.5% divergence among duplicated genomes | 30 | 24.16 | 11.75 | 0.486 | 1,197 | 9,818 | 712.9 | 700.4 | 0.982 |
| 1% divergence | 30 | 24.16 | 13.09 | 0.542 | 6,699 | 1,954 | 713.0 | 696.8 | 0.977 |
| 5% divergence | 30 | 24.16 | 22.92 | 0.949 | 4,719 | 4,857 | 714.4 | 700.3 | 0.980 |

The least relative deviation from the true genome size overall was found for the N_clean_/*λ* estimator (mean deviation 0.00017, range 0.00003-0.00056), followed by N_clean_/*m* (0.054, 0.0169 - 0.111), N_bm_/*m* (0.094, 0.014-0.209) and N_bm_/*λ* (0.112, 0.003-0.224, Figure 1). There was a tendency for the method to perform better with higher coverage, mainly due to the smaller relative deviation of *m* from *λ* at higher coverage. Given the rather minor differences in contiguity among genome assemblies reconstructed from different coverages, this factor had only a minor role for the precision of the genomes size estimates (Table 3, Figure 1).

**Figure 1.**
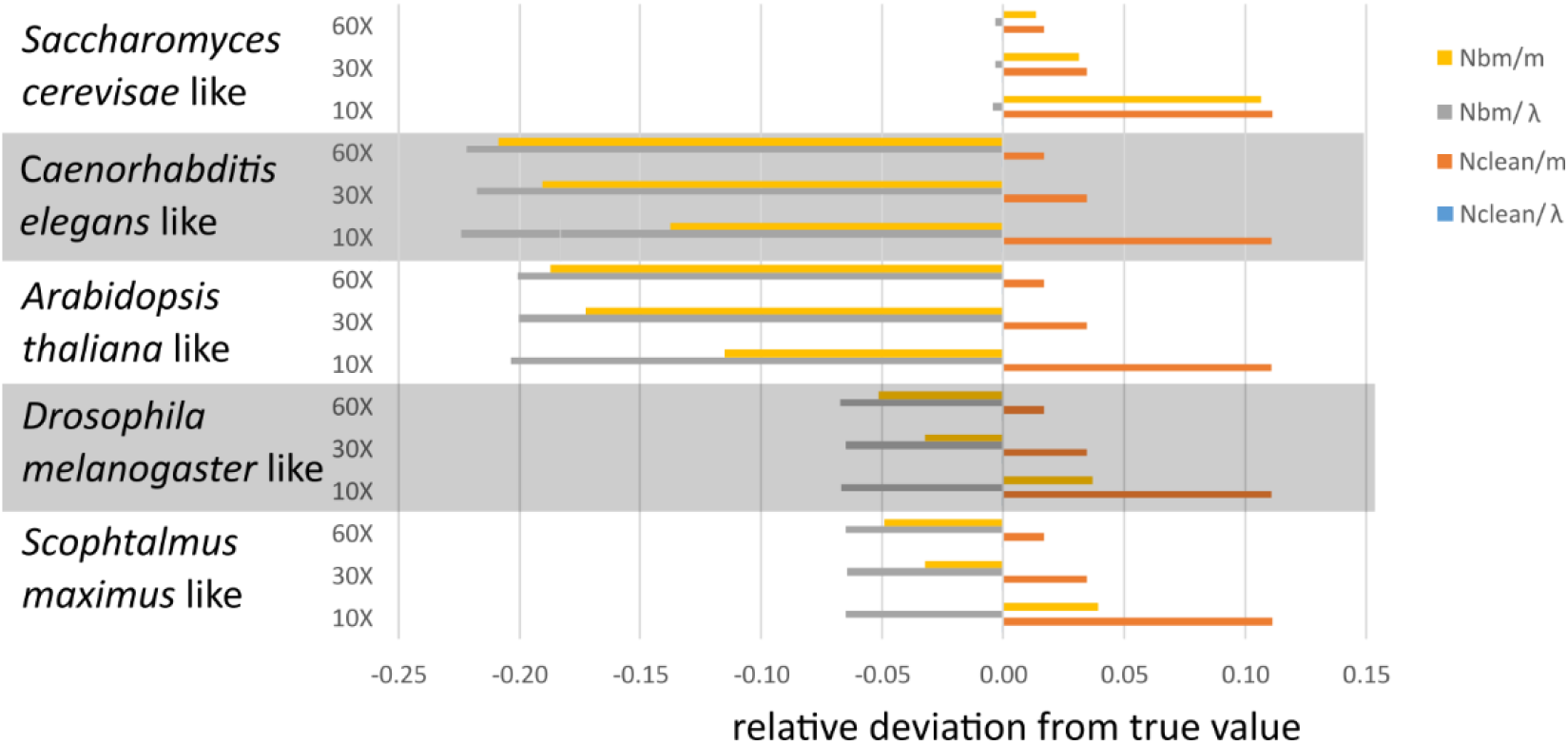
Relative deviations of genome size estimators from true values for different simulated genomes and simulated coverages. The deviations of N_clean_/*λ* (blue) from the true value are so small that they are not visible on the scale. The raw data table to this figure can be found in the Supplemental Table 1.

The genome size estimates from simulated genomes with varying proportions of divergent heterozygous sites all yielded the same estimates (Supplemental Table 1). As can be seen in the respective coverage distributions, the only difference between the simulations was a second, lower peak at about half the expected coverage that grew with increasing amount of heterozygous regions. The position of the true peak remained unaffected (Figure 3a).

**Figure 3.**
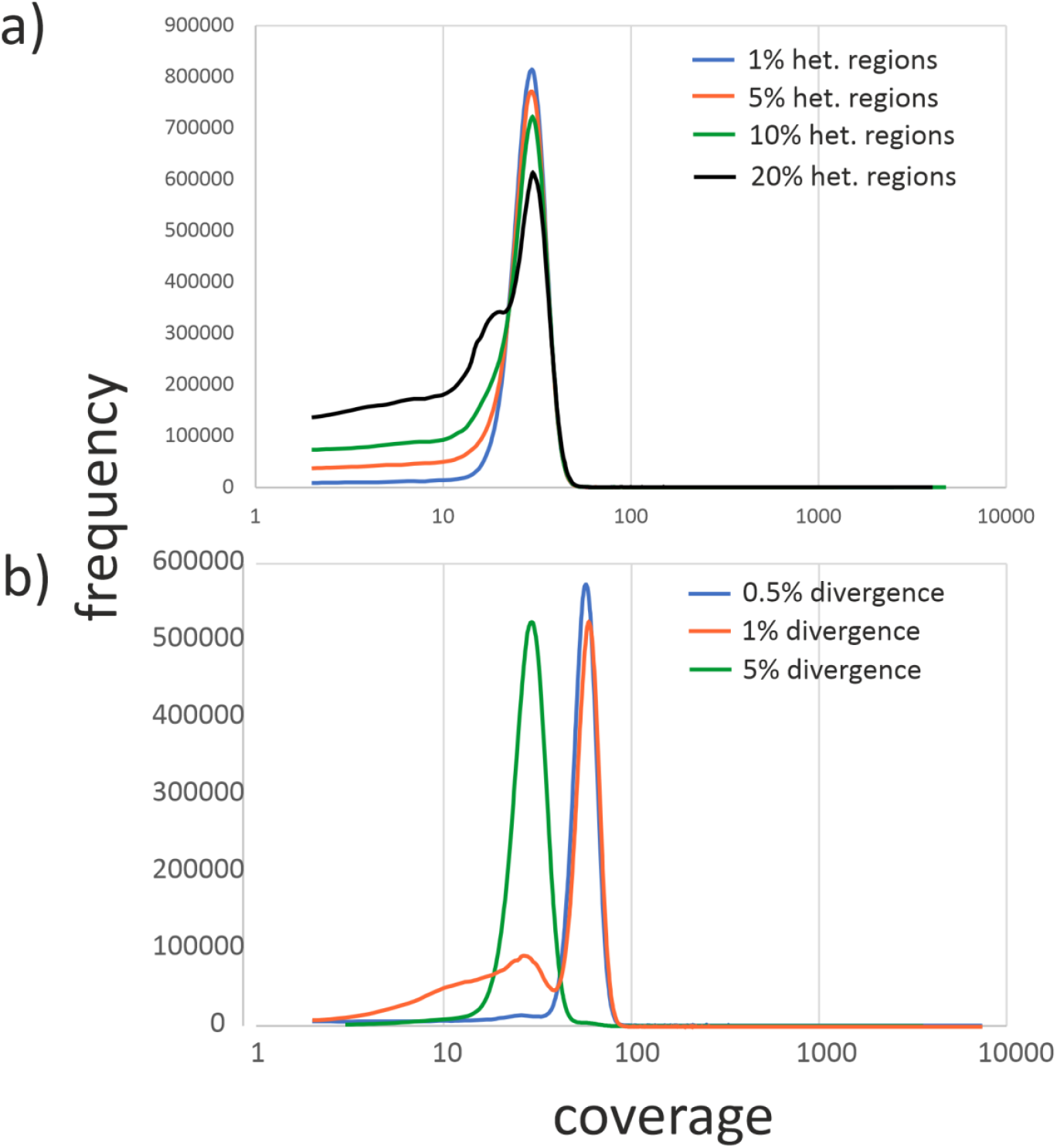
Coverage distributions for divergent heterozygous and tetraploid genomes. All distributions shown are based on the *Saccharomyces_cerevisae*_like genome. a) Coverage distributions for 0%, 5% 10% and 20% of divergent heterozygous regions. b) Coverage distributions for tetrapoid genomes with 0.5%, 1% and 5% divergence among the duplicated genomes. Please note the logarithmic scale of the x-axes.

Assembly of a tetraploid *Saccharomyces cerevisae*-like genome with the two lowest divergences between the duplicated genomes (0.5% and 1%) resulted in the reconstruction of approximately a single haploid genome, respectively (assemblies of lengths 1.18 Mb and 1.31 Mb, Supplemental Table 1). Therefore, the highest observed coverages for these simulations were both 59 and the *λ* estimates close to 60 (Supplemental Table 1, Figure 3b). Consequently, the genome size estimates were close to the haploid length. However, with divergence 1%, a second peak with maximum 28, respectively *λ* 28.9 emerged (Figure 3b, Supplemental Table 1). Using this peak yielded estimates that were much closer to the truth (relative deviations between 0.005 and 0.06, depending on estimator). With 5% divergence, the duplicated genomes were almost fully resolved in the assembly and, hence, the peak at the true coverage and therefore the genome size estimates not further than 0.03 from the truth (Figure 3b, Supplemental Table 1).

### Empirical data

Ordinary Least Squares Regression of 1C flow-cytometry estimates against the estimates derived from the coverage approach yielded an excellent fit (r^2^ = 0.998, p = 2.2 × 10^−34^). Removing the outlier estimate for the crayfish genome did not change the result markedly (r^2^ = 0.958, p = 2.1 × 10^−17^). The estimated slope was with 0.996+/-0.043 (s.e.) very close to unity. The fit of the respective k-mer based estimates to the flow cytometry data was equally good (r^2^ = 0.996, p = 1.1 × 10^−26^), however, the slope of 0.585 +/-0.007 (s.e.) suggested a systematically lower k-mer estimate (Figure 2). The estimated slopes were significantly different from each other (t = 9.43, d.f. = 44, p < 1 × 10^−6^).

**Figure 5.**
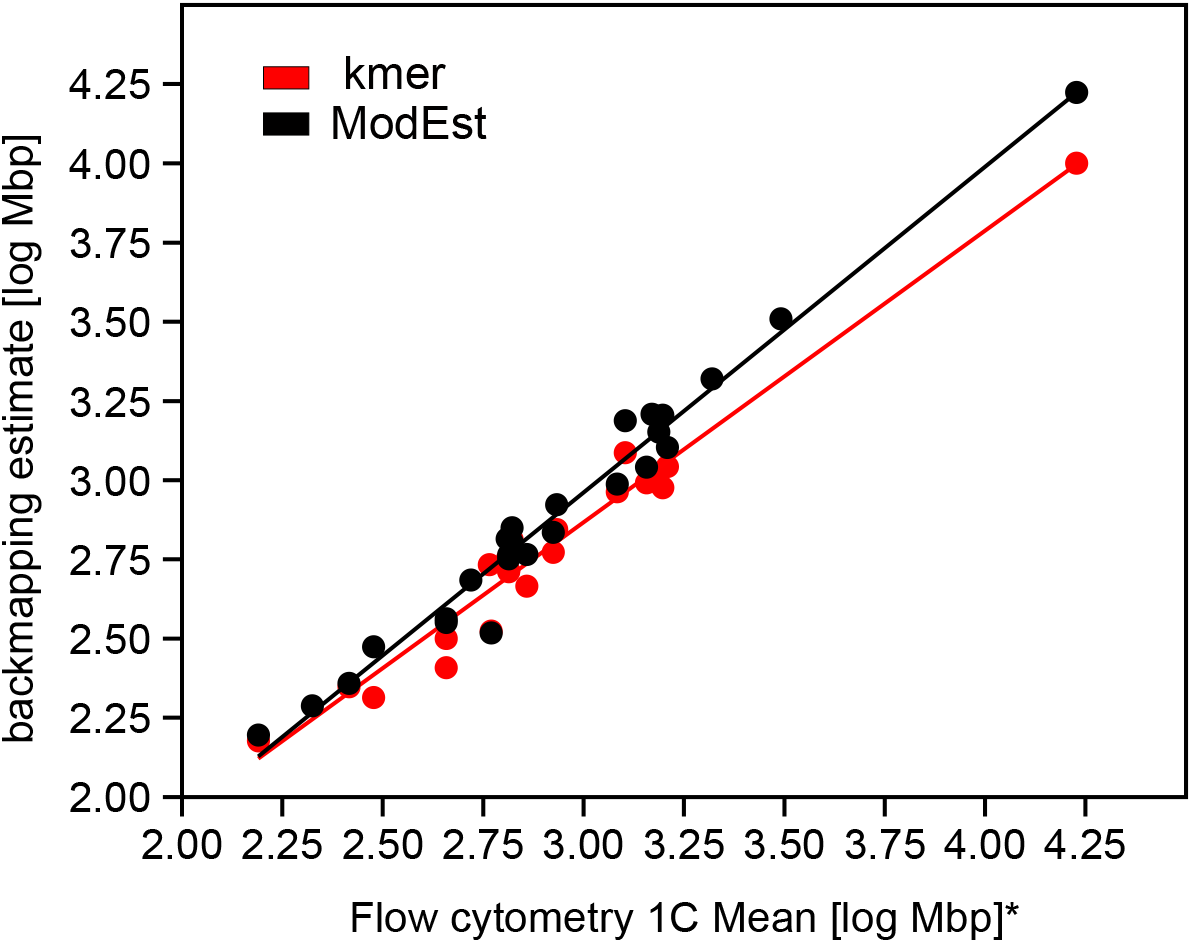
Ordinary least square regression for N_bm_/m (black) and k-mer based (red) genome size estimates on 1C flow-cytometry estimates derived from the same individuals, respectively. For better graphical representation, estimates were log transformed. Both regressions were highly significant (p < 0.0001). The N_bm_/m estimates fit as well (r^2^ = 0.998) than their k-mer based counterparts (r^2^ = 0.996). The slopes (0.995 for N_bm_/m and 0.59 for k-mer based) were significantly different.

## Discussion

As long as reliable whole chromosome sequencing is technically not yet feasible and thus the true genome size is not known, genome size estimation of *de novo* sequenced genomes will be a necessary and important part of biodiversity genomics. We presented here with ModEst a fast, easy- to-use and precise method for genome size estimation from NGS sequencing data. We have shown that the method works for a wide range of genome sizes. The method could become standard part of the genome assembly process, because it relies on data that is routinely collected. Albeit our method is not the first to propose the use of sequencing, respectively mapping statistics (Pflug *et al*. 2020; Pucker 2019), it requires less assumptions and much less bioinformatic effort than previously suggested approaches. The method does, admittedly, not solve the problem how much sequence information should be produced in the first place if there is absolutely no *a priori* information on the expected genome size of the target organism. However, very low modal coverages obtained with the method indicate that sequencing efforts should be increased.

To evaluate the performance of our method and the factors influencing it, we performed a simulation study. We simulated five different genomes with the characteristics and genome sizes typical for various eukaryotic taxa. We could show that the precision of the estimate is largely independent from the contiguity and quality of the underlying genome assembly as long as most sequence elements in the genome are represented in the assembly draft. This finding was confirmed with the empirical samples, where e.g. the size estimate for giant genome of the crayfish *Astacus astacus* was gained from a very preliminary, highly discontinuous assembly with poor N50, which nevertheless yielded excellent concordance with the flowcytometry estimates (Table 2). This makes the method particularly suitable to obtain a reliable genome size estimate early in the assembly process and, if necessary, adjust the sequencing strategy. But also genome skimming projects (Dodsworth 2015) with low coverages could profit from the proposed method, as long as the obtained coverage is at least in the order of 2-5X. The simulations have further shown that divergent heterozygote stretches do not compromise the result of the genome size estimation.

The accuracy of genome size estimates of simulated tetraploid organisms depended strongly on the degree of divergence between the genome copies. When the divergence was low (0.5%), the assembly of the duplicated was almost completely collapsed and consequently the modal coverage twice as high as the true coverage. However, already with 1% sequence divergence between the duplicated genomes, an additional peak close to the true value of 30 was observed. For 5% sequence divergence and higher (not shown), the assembly more or less fully resolved the duplicated genomes and the highest peak was identical to the true coverage. This stressed that multimodal coverage distributions point to issues with the assembly and should always be carefully investigated.

Nevertheless, if the ploidy of the organism is known, reliable estimates of the genome size can be gained even for recent polyploidisation events with our method as well.

The simulation study relied on simulated short reads as obtained e.g. by the widespread Illumina-platform. However, several included empirical examples (e.g. Chueca *et al*. 2021a) suggested that estimating the bases sequenced from the target genome with PacBio long reads worked equally well. In principle, as long as the assumption of random sequencing of bases from the genome is fulfilled, every sequencing platform should yield reliable estimates. For mixed assemblies, however, it is advisable to use only one sort of data (preferably the one with the higher number of sequenced bases, see below), because the underlying coverage distributions are usually different.

We proposed four slightly different estimators of genome size. Simulations indicated that, as expected, the N_clean_/*λ* estimator yielded by far the best results, in practice largely independent of coverage or assembly quality. However, since we gained the reads from simulated genomes, they were by definition free of contaminations, i.e. reads from other organisms or other (e.g. organellar) genomes. Whether N_clean,_ the number of bases sequenced after cleaning and trimming, is reasonable for empirical estimations depends thus on the confidence with regard to the amount of residual contamination in the data set.

For the alternative, using the number of back-mapped reads, N_bm_, as an estimator of the bases sequenced, precision depended strongly on the completeness of the genome assembly in terms of presence of all sequence elements, regardless of their copy-number. This seemed reasonable: if all repeat classes and complex regions are represented in the genome draft, all reads will find a place they can map to. If the confidence is high that N_clean_ is correct, the ratio N_bm_/N_clean_ would be a good indicator of genome completeness in this sense.

We have shown that the *λ* parameter of the underlying true Poisson distribution of base coverage is readily and reliably found by ML estimation, if we truncate the data to a small window around the modal value of the coverage distribution. Moreover, because the modal value of a Poisson distribution cannot deviate more than 1 from *λ*, the relative error from using m instead of *λ* decreases with increasing coverage. Most genome sequencing projects use coverages of several dozen X for at least one technique where the difference becomes marginal. Estimating genome size from low coverage e.g. of genome-skimming projects, however, should entail proper estimation of *λ*.

Comparison of genome size estimates obtained with our sequencing coverage method to empirical data from flow cytometry obtained from the same individual achieved very good agreement, regardless of genome size. The regression slope of close to 1 indicated that the estimates obtained with our method can be used interchangeably with those from flow-cytometry. This allows researchers to gather reliable and comparable genome size estimates for species where fresh material is difficult or impossible to obtain or access to flow-cytometry equipment is lacking.

While the k-mer based estimates available were almost as consistent as those obtained from sequencing coverage, they were not as precise. The k-mer approach consistently underestimated the true size by more than one third. By their very nature, k-mer approaches estimate rather the content of high complexity regions (Lipovský *et al*. 2017). It will be therefore interesting to see whether the observed taxon-independent relationship of approximately 2/3 complexity regions to 1/3 repeat regions as found here mainly for animal species will hold true for more genomes. The work of Novák et al. (2020) also showed an almost constant, albeit higher proportion of repetitive regions for plant genomes with sizes up to 10 Gb.Above this size, the relative proportion of repeats declined. Obtaining more reliable genome sizes from a broad taxon range will allow to infer which processes are driving these patterns to which the proposed ModEst method can contribute.

## Acknowledgements

We thank our LOEWE-TBG colleagues for giving us early access to their assembled genomes.

## Data Accessibility Statement

All genomes specifically simulated for this publication will be made available via Dryad.

## Supplement

R-code for estimating lambda from a truncated Poisson distribution

library(fitdistrplus)

library(truncdist)

library(splitstackshape)

#transform Qualimap output to R-object

obj <-read.table(“coverage_histogram.txt”, header = TRUE)

obj <-expandRows(obj, “freq”)

obj <-as.vector(obj$freq)

summary(obj)

#define function for mode

mode <-function(obj) {uniqv <-unique(obj) uniqv[which.max(tabulate(match(obj, uniqv)))]}

min <-mode – 5

max <-mode + 5

dtruncated_poisson <-function(x, lambda) {dtrunc(x, “pois”, a=min, b=max, lambda=lambda)}

ptruncated_poisson <-function(q, lambda) {ptrunc(q, “pois”, a=min, b=max, lambda=lambda)}

fitdist(obj, “pois”, start = list(lambda = mode))

**Supplemental Table 1.** Genome size estimates and deviations from true value for the simulated genomes.

| simulated genome | coverage | estimated | modal | $N_{\text{clean}}/\lambda$ | deviation | $N_{\text{clean}}/m$ | deviation | $N_{\text{bm}}/\lambda$ | deviation | $N_{\text{bm}}/m$ | $N_{\text{bm}}/m$ |
| --- | --- | --- | --- | --- | --- | --- | --- | --- | --- | --- | --- |
| | | $\lambda$ | coverage | | $N_{\text{clean}}/\lambda$ | | $N_{\text{clean}}/m$ | | $N_{\text{bm}}/\lambda$ | | |
|  |  |  | m |  |  |  |  |  |  |  |  |
| **Saccharomyces_cerevisae_like** | 10X | 9.995 | 9 | 12.08 | -0.00044 | 13.42 | 0.11109 | 11.30 | -0.06500 | 12.56 | 0.03933 |
|  | 30X | 29.994 | 29 | 12.08 | -0.00015 | 12.50 | 0.03446 | 11.30 | -0.06447 | 11.69 | -0.03209 |
|  | 60X | 59.998 | 59 | 12.08 | -0.00003 | 12.29 | 0.01693 | 11.29 | -0.06505 | 11.49 | -0.04920 |
| **Caenorhabditis_elegans_like** | 10X | 9.996 | 9 | 100.04 | -0.00010 | 111.15 | 0.11103 | 93.37 | -0.06678 | 103.74 | 0.03694 |
|  | 30X | 29.995 | 29 | 100.03 | -0.00019 | 103.49 | 0.03441 | 93.56 | -0.06487 | 96.79 | -0.03251 |
|  | 60X | 59.994 | 59 | 100.04 | -0.00003 | 101.73 | 0.01688 | 93.31 | -0.06736 | 94.88 | -0.05159 |
| **Arabidopsis_thaliana_like** | 10X | 9.993 | 9 | 120.04 | -0.00038 | 133.42 | 0.11102 | 95.63 | -0.20365 | 106.29 | -0.11490 |
|  | 30X | 29.996 | 29 | 120.07 | -0.00016 | 124.22 | 0.03440 | 96.02 | -0.20037 | 99.34 | -0.17273 |
|  | 60X | 59.992 | 59 | 120.08 | -0.00005 | 122.11 | 0.01687 | 95.98 | -0.20075 | 97.60 | -0.18723 |
| **Drosophila_melanogaster_like** | 10X | 9.998 | 9 | 143.98 | -0.00056 | 160.06 | 0.11100 | 111.75 | -0.22432 | 124.22 | -0.13773 |
|  | 30X | 29.992 | 29 | 144.05 | -0.00009 | 149.02 | 0.03438 | 112.72 | -0.21756 | 116.61 | -0.19059 |
|  | 60X | 59.988 | 59 | 144.06 | -0.00003 | 146.49 | 0.01685 | 112.09 | -0.22198 | 113.98 | -0.20884 |
| **Scophthalmus_maximus_like** | 10X | 9.996 | 9 | 523.93 | -0.00027 | 582.29 | 0.11110 | 521.90 | -0.00414 | 580.04 | 0.10680 |
|  | 30X | 29.995 | 29 | 524.05 | -0.00004 | 542.13 | 0.03448 | 522.34 | -0.00330 | 540.37 | 0.03111 |
|  | 60X | 59.998 | 59 | 524.03 | -0.00008 | 532.95 | 0.01694 | 522.27 | -0.00344 | 531.16 | 0.01353 |
| **Saccharomyces_cerevisae_like 1% divergent heterozygous regions** | 30X | 29.999 | 30 | 11.87 | -0,01747 | 11.87 | -0,01749 | 11.74 | -0,02827 | 11.74 | -0,02829 |
| **5% divergent heterozygous regions** | 30X | 29.999 | 30 | 11.80 | -0,02321 | 11.80 | -0,02324 | 11.68 | -0,03286 | 11.68 | -0,03288 |
| **10% divergent heterozygous regions** | 30X | 29.999 | 30 | 11.71 | -0,03038 | 11.71 | -0,03040 | 11.57 | -0,04228 | 11.57 | -0,04230 |
| **20% divergent heterozygous regions** | 30X | 29.999 | 30 | 11.54 | -0,04504 | 11.54 | -0,04507 | 11.40 | -0,05666 | 11.40 | -0,05668 |
| **Tetraploid**<br>**Saccharomyces_cerevisae_like 0.5%**<br>**divergence among duplicated**<br>**genomes** | 30X | 59.999 | 59 | 11.88 | -0.50819 | 12.08 | -0.49986 | 11.67 | -0.51685 | 11.87 | -0.50867 |
| **1% divergence** | 30X | 59.988 | 59 | 11.89 | -0.50804 | 12.08 | -0.49980 | 11.62 | -0.51924 | 11.81 | -0.51119 |
| **1% divergence, correct peak** | 30X | 28.986 | 28 | 24.60 | 0.01815 | 25.46 | 0.05398 | 24.04 | -0.00502 | 24.88 | 0.03000 |
| **5% divergence** | 30X | 29.999 | 29 | 23.81 | -0.01431 | 24.63 | 0.01965 | 23.34 | -0.03376 | 24.15 | 0.00047 |

